# The Function of the COPII Gene Paralogs SEC23A and SEC23B Are Interchangeable *In Vivo*

**DOI:** 10.1101/215160

**Authors:** Rami Khoriaty, Geoffrey Hesketh, Amélie Bernard, Angela C. Weyand, Dattatreya Mellacheruvu, Guojing Zhu, Mark J. Hoenerhoff, Beth McGee, Lesley Everett, Elizabeth J. Adams, Bin Zhang, Thomas Saunders, Alexey Nesvizhskii, Daniel J. Klionsky, Jordan A. Shavit, Anne-Claude Gingras, David Ginsburg

## Abstract

SEC23 is a core component of the coat protein-complex II (COPII)-coated vesicle, which mediates transport of secretory proteins from the endoplasmic reticulum (ER) to the Golgi^1-3^. Mammals express 2 paralogs for SEC23 (SEC23A and SEC23B). Though the *SEC23* gene duplication dates back >500 million years, both SEC23’s are ~85% identical at the amino acid sequence level. In humans, deficiency for SEC23A or SEC23B results in cranio-lenticulo-sutural dysplasia^4^ or congenital dyserythropoietic anemia type II (CDAII), respectively^5^. The disparate human syndromes and reports of secretory cargos with apparent paralog-specific dependence^6,7^, suggest unique functions for the two SEC23 paralogs. Here we show indistinguishable intracellular interactomes for human SEC23A and SEC23B, complementation of yeast SEC23 by both human and murine SEC23A/B paralogs, and the rescue of lethality resulting from *Sec23b* disruption in zebrafish by a *Sec23a*-expressing transgene. Finally, we demonstrate that the *Sec23a* coding sequence inserted into the endogenous murine *Sec23b* locus fully rescues the mortality and severe pancreatic phenotype previously reported with SEC23B-deficiency in the mouse^8-10^. Taken together, these data indicate that the disparate phenotypes of SEC23A and SEC23B deficiency likely result from evolutionary shifts in gene expression program rather than differences in protein function, a paradigm likely applicable to other sets of paralogous genes. These findings also suggest the potential for increased expression of SEC23A as a novel therapeutic approach to the treatment of CDAII, with potential relevance to other disorders due to mutations in paralogous genes.

---

Recruitment of SAR1-SEC23-SEC24 to the cytoplasmic face of the ER membrane forms the inner coat of the COPII vesicle. Interaction of SEC23 with each of the 4 SEC24 mammalian paralogs (SEC24A-D) provides a potential for functional differentiation between SEC23A and SEC23B, although we detected no differences in paralog specificity by co-immunoprecipitation (Extended Data Fig. 1). To comprehensively characterize the full set of interacting proteins for each SEC23 paralog, we performed 2 independent “BioID” (proximity-dependent biotinylation) experiments in HEK293 cells equivalently expressing either BirA*-tagged SEC23A, SEC23B, or GFP control (Fig. 1a, b). Comparison of the SEC23A to SEC23B interactomes revealed a remarkably high correlation (r^2^ = 0.96 and 0.95) in two independent experiments (Fig. 1c,d and Extended Figure 2 a-c), equivalent to the correlation between biological replicates for SEC23A (r^2^ = 0.98, Fig. 1e) or SEC23B (r^2^ = 0.98, Fig. 1f). Thus, the interactomes of SEC23A and SEC23B in HEK293T cells are indistinguishable by BioID.

**Figure 1.**
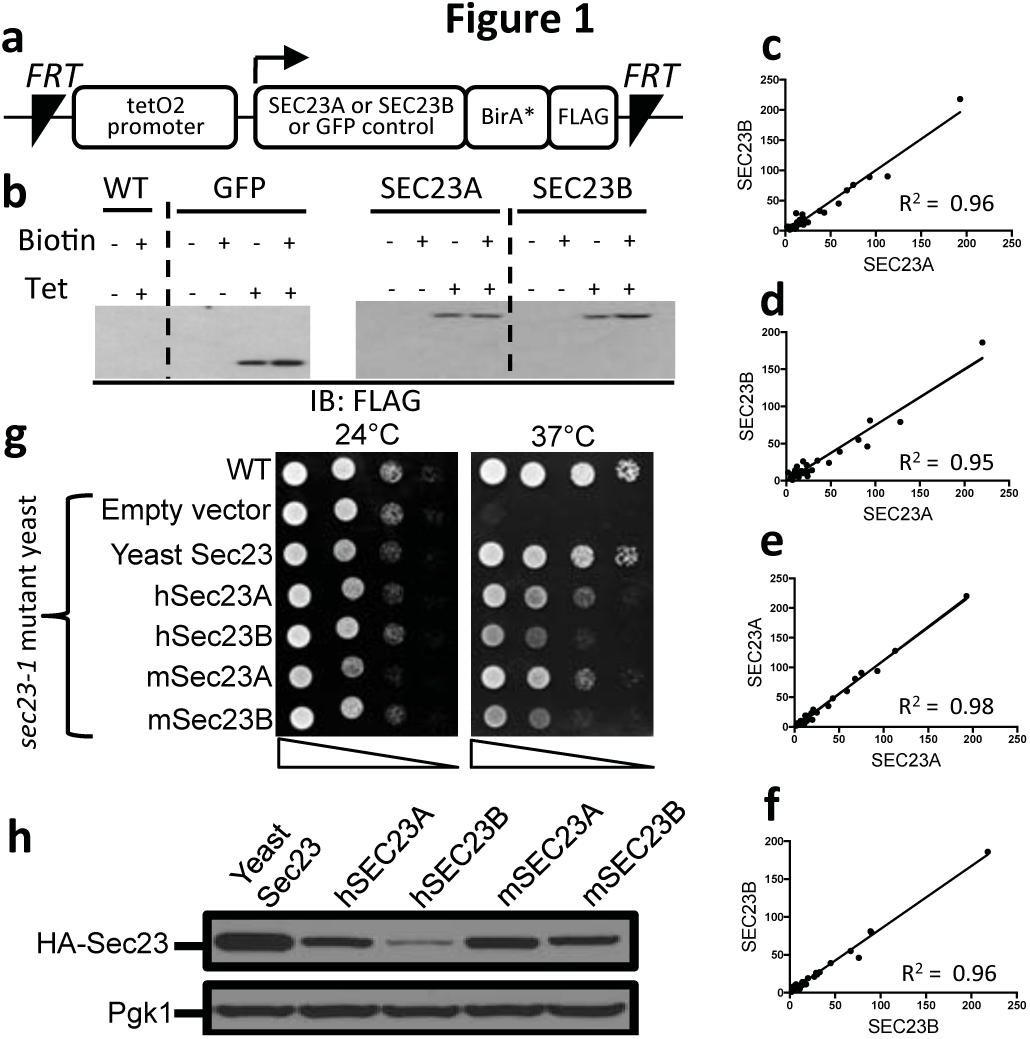
SEC23A and SEC23B exhibit indistinguishable interactomes in HEK293 cells and complement the yeast SEC23 protein. **a**, TREX HEK293 cells expressing SEC23A or SEC23B with a BirA* and FLAG tags. **b**, FLAG immunoblotting following tetracycline induction. **c-f**, SEC23A and SEC23B interactomes were each determined in 2 independent experiments. Pairwise comparisons of spectral counts for a given protein are shown across paralogs (**c,d**) or between replicas for the same paralog (**e,f**). **g**, Growth of temperature-sensitive *sec23-1* mutant yeast expressing HA-tagged human (h) or mouse (m) SEC23 paralogs (or yeast SEC23) from the *ZEO1* promoter at the permissive and restrictive temperatures. **h**, Immunoblotting for HA detects relative expression for each construct.

**Figure 2.**
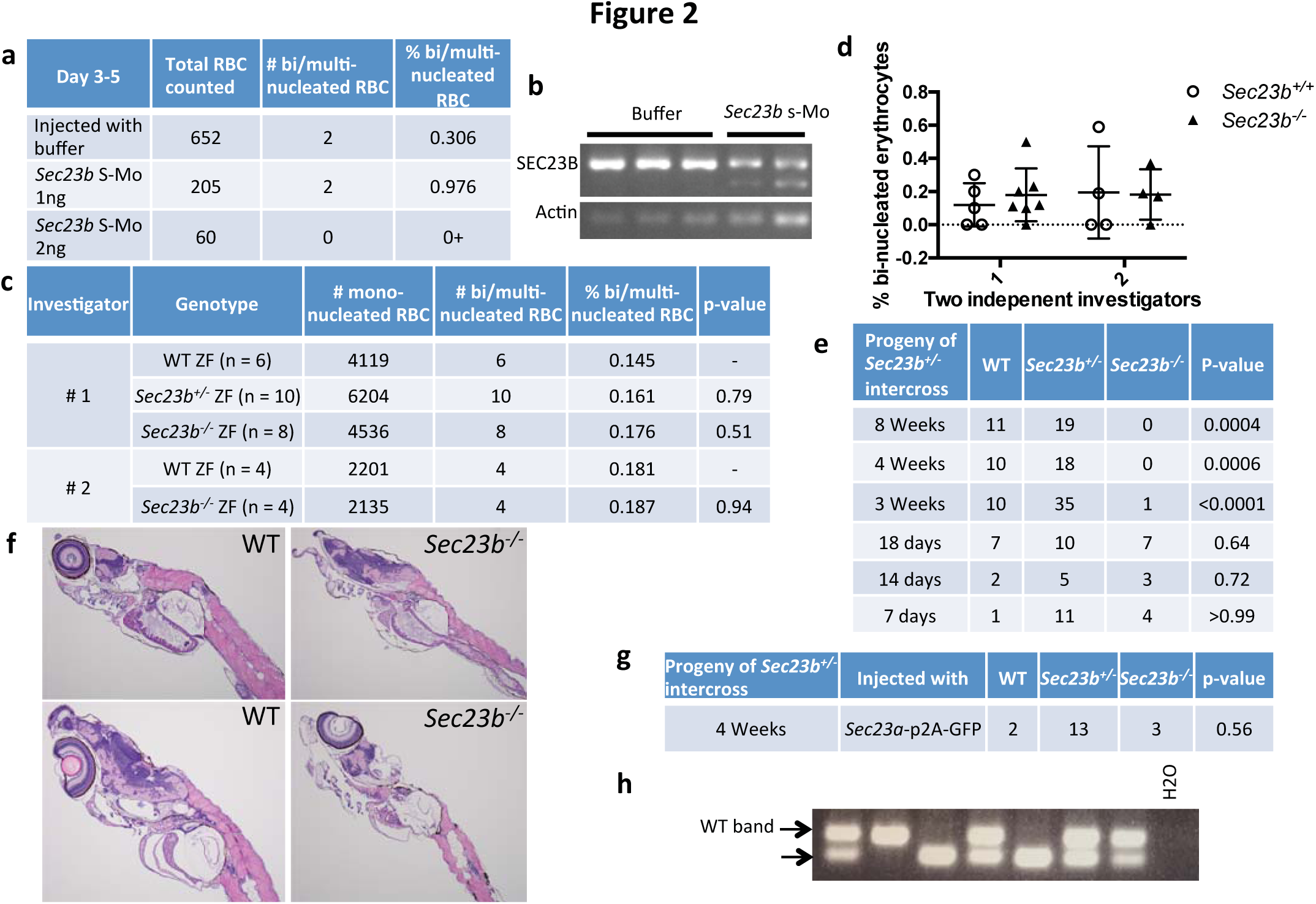
Mortality in SEC23B-deficient zebrafish (ZF) between 18-21 days, rescued by SEC23A. **a**, Number of peripheral blood binucleated erythrocytes and **b**, RT-PCR for *Sec23b* following injection of 1 or 2 ng *Sec23b* morpholino. **c,d**, Peripheral blood binucleated erythrocyte counts (2 independent investigators) from *Sec23b*^*−/−*^ ZF (CRISPR/Cas9-induced 53bp deletion). **e**, *Sec23b*^*−/−*^ ZF die between 18-21 days. **f**, H&E stains at day 16. **g,h**, One-cell stage embryos from *Sec23b*^*+/−*^ ZF intercrosses were injected with a *Sec23a* transgene. **h**, Genotyping at 4 weeks of age demonstrates rescue of *Sec23b*^*−/−*^ mortality by SEC23A.

Human SEC23A but not human SEC23B was previously reported to complement the yeast Sec23 protein^11^, a puzzling finding in light of the above evidence for indistinguishable SEC23A/B interactomes in mammalian cells. We thus re-tested complementation in a temperature sensitive *sec23-1* mutant yeast strain using both human (h) and mouse (m) SEC23A and B sequences. Indeed, all 4 mammalian sequences (hSEC23A, hSEC23B, mSEC23A and mSEC23B) rescued growth of the temperature sensitive *sec23-1* mutant yeast at the restrictive temperature (Fig. 1g, h). The discrepancy between these findings and the previous report^11^ is likely explained by poor expression of hSEC23B in yeast compared to hSEC23A (Extended Data Fig. 3 a-d), potentially due to differences in codon usage or protein stability.

**Figure 3.**
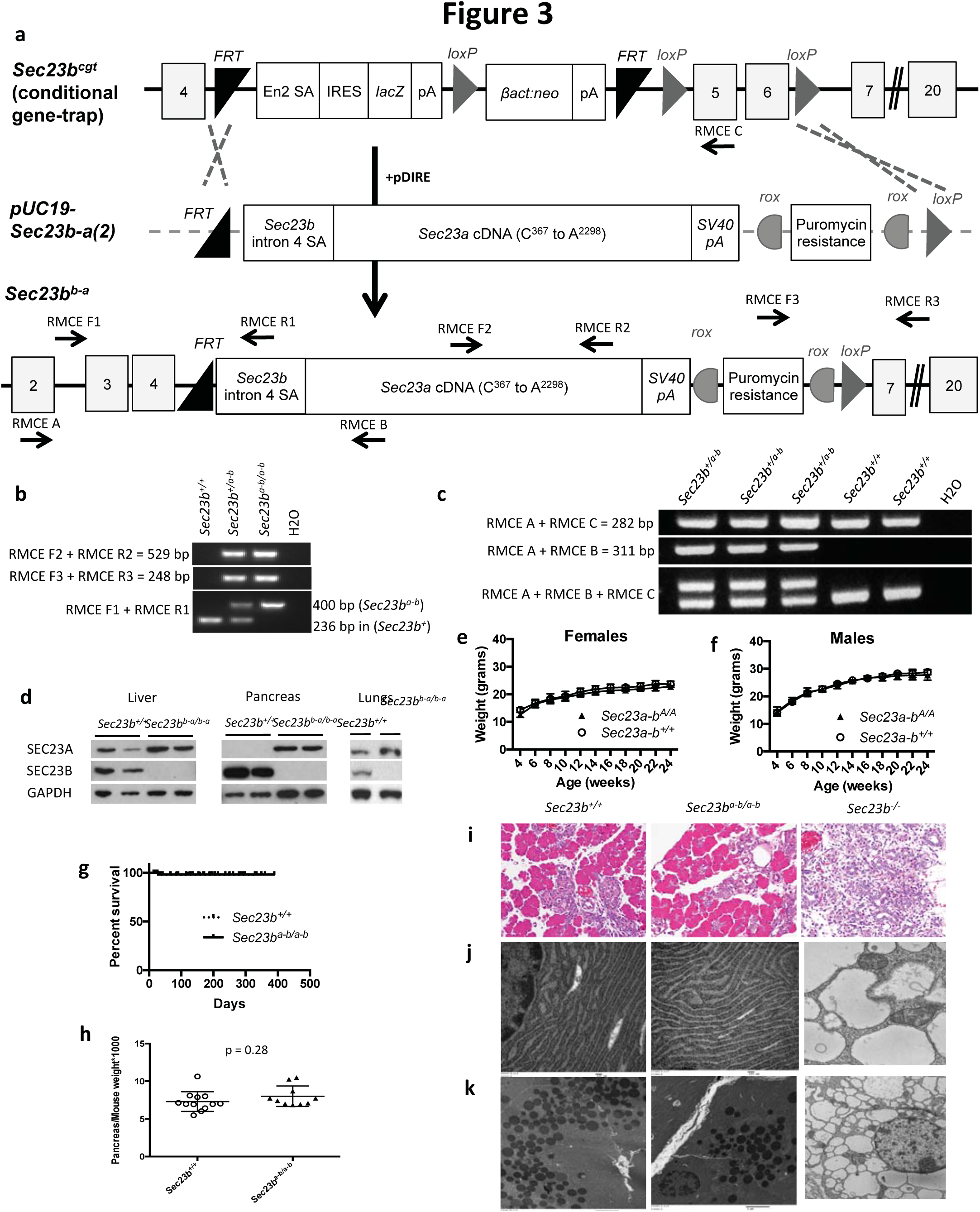
Mice expressing *Sec23a* from the *Sec23b* locus (*Sec23b*^*b-a/b-a*^) exhibit normal growth, pancreas histology, and absence of pancreatic ER dilation. **a**, Design of the replacement vector pUC19-Sec23b-a(2). Arrows indicate locations of the genotyping primers. **b**, PCR demonstrates the expected band sizes for correct *Sec23b*^*b-a*^ 5’ and 3’ insertions. **c**, RT-PCR identifies the expected splicing of *Sec23b* exon4 into the *Sec23a* cDNA. **d**, Western blot for SEC23A and SEC23B in tissues from *Sec23b*^*b-a/b-a*^ and WT mice. **e,f** Growth and **g**, survival of *Sec23b*^*b-a/b-a*^ mice. **h, i**, Weight and histology of *Sec23b*^*b-a/b-a*^ pancreas tissues. **j**, Appearance of ER and **k**, zymogen granules in *Sec23b*^*b-a/b-a*^ pancreas tissues by electron microscopy.

SEC23B deficiency in humans results in CDAII, a disease characterized by anemia and increased bi-nucleated erythroid precursors^2^, with *Sec23b* knockdown in ZF previously reported to similarly result in an increased percentage of circulating bi-nucleated erythrocytes^5^. In contrast, SEC23B deficiency in mice results in perinatal mortality and massive pancreatic degeneration but no findings comparable to CDAII ^8-10,12^. To address these discordant phenotypes across species, we repeated the injection of one-cell stage ZF embryos with the identical *Sec23b* splice-blocking Mo reported by Schwarz et al^5^. In contrast to the previous report, we observed no increase in circulating bi-nucleated erythroid cells from *Sec23b* knockdown (Fig. 2a, and Extended Data Table 2). To exclude an effect of residual *Sec23b* expression (Fig. 2b), ZF heterozygous for a 53-bp deletion in exon 5 of *Sec23b* (*Sec23b*^*+/−*^) were generated by CRISPR/Cas9 genome editing. No difference in the percentage of circulating bi-nucleated erythrocytes was noted at day 16 between *Sec23b*^*−/−*^ ZF compared to WT and *Sec23b*^*+/−*^ ZF clutch mates (Fig. 2d, e). Interestingly, however, none of the *Sec23b*^*−/−*^ ZF survived beyond 3 weeks of age, despite absence of any gross or histologic abnormalities following pathology evaluation (Fig. 2c, f).

To determine if SEC23A could rescue the early mortality of SEC23B-deficient ZF, one-cell stage embryos generated from *Sec23b*^*+/−*^ intercrosses were injected with a SEC23A or GFP transgene (Extended Data Fig. 4a). Genotyping of transgene-expressing ZF demonstrated that SEC23A rescues the mortality of SEC23B-deficiency in ZF (Fig. 2g, h and Extended Data Fig. 4 b-d).

The ZF results suggest at least partial overlap in function between SEC23A and SEC23B. To characterize the capacity of the SEC23A protein to replace SEC23B in mammals *in vivo*, we substituted SEC23A coding sequences for SEC23B (at the endogenous *Sec23b* genomic locus). Although efforts to engineer this allele by direct oocyte injection were unsuccessful (Extended Data Fig. 5), embryonic stem (ES) cells carrying the Sec23b-a allele were generated as shown in Fig. 3a, and used to obtain mice carrying the *Sec23b*^*b-a*^ allele. *Sec23b*^*+/b-a*^ mice (Fig. 3b, c) were observed at the expected Mendelian ratio in N2 progeny (p = 0.47, Table 1a). *Sec23b*^*+/b-a*^ mice were indistinguishable from their WT control littermates, exhibiting normal growth, fertility and gross appearance, and no abnormalities on standard pathology evaluation.

**Table 1.** Mouse crosses to determine if SEC23B deficient mice expressing *Sec23a* from the *Sec23b* locus are viable. As shown in (A-B), mice expressing *Sec23a* from the *Sec23b* locus are viable at weaning. To test the capacity of the *Sec23b*^*b-a*^ allele to rescue the lethality of mice homozygous for an inactivated endogenous *Sec23a* allele (*Sec23a*^*−/−*^ mice), *Sec23b*^*+/a-b*^ mice were crossed with the *Sec23a*^*+/−*^ mice. Although, the expected number of *Sec23b*^*+/b-a*^ *Sec23a*^*+/−*^ mice from a *Sec23b*^*+/b-a*^ *Sec23a*^*+/−*^ x *Sec23a*^*+/−*^ cross were present at weaning (C), no *Sec23b*^*+/b-a*^ *Sec23a*^*−/−*^ mice were observed at weaning or at E13.5 (D). Thus *Sec23b*^*b-a*^ fails to complement loss of Sec23a expression in *Sec23a*^*−/−*^ mice. * p-value calculated for *Sec23b*^*+/b-a*^versus wild type mice. A p-value calculated for for *Sec23b*^*b-a/b-a*^ versus all other genotypes. ^#^ p-value calculated for *Sec23a*^*+/−*^ *Sec23b*^*+/b-a*^ versus all other genotypes. ^¥^ p-value calculated for *Sec23a*^*−/−*^ *Sec23b*^*+/b-a*^ mice versus all other genotypes.

|  |  |  |  |  |  |  |
| --- | --- | --- | --- | --- | --- | --- |
| **A. Genotypes** | *Sec23b*<sup>+/+</sup> | *Sec23b*<sup>+/b-a</sup> | - | - | - | **p-value\*** |
| *Sec23b*<sup>+/b-a</sup> x *Sec23b*<sup>+/+</sup><br>**Expected ratios** | **50%** | **50%** | - | - | - |  |
| Observed at weaning % (n=324) | 47.8% (155) | 52.2% (169) | - | - | - | 0.47 |
| **B. Genotypes** | *Sec23b*<sup>+/+</sup> | *Sec23b*<sup>+/b-a</sup> | *Sec23b*<sup>b-a/b-a</sup> | - | - | **p-value^** |
| *Sec23b*<sup>+/b-a</sup> x *Sec23b*<sup>+/b-a</sup><br>**Expected ratios** | **25%** | **50%** | **25%** | - | - |  |
| Observed at weaning % (n=287) | 23.4% (67) | 50.5% (145) | 26.1% (75) | - | - | 0.68 |
| **C. Genotypes** | *Sec23a*<sup>+/+</sup><br>*Sec23b*<sup>+/+</sup> | *Sec23a*<sup>+/+</sup><br>*Sec23b*<sup>+/b-a</sup> | *Sec23a*<sup>+/-</sup><br>*Sec23b*<sup>+/+</sup> | *Sec23a*<sup>+/-</sup><br>*Sec23b*<sup>+/b-a</sup> |  | **p-value#** |
| *Sec23a*<sup>+/-</sup> x *Sec23b*<sup>+/b-a</sup><br>**Expected ratios** | **25%** | **25%** | **25%** | **25%** | - |  |
| Observed at weaning % (n=79) | 29.1% (23) | 21.5% (17) | 29.1% (23) | 20.3% (16) | - | 0.37 |
| **D. Genotypes** | *Sec23a*<sup>+/+</sup><br>*Sec23b*<sup>+/+</sup> | *Sec23a*<sup>+/+</sup><br>*Sec23b*<sup>+/b-a</sup> | *Sec23a*<sup>+/-</sup><br>*Sec23b*<sup>+/+</sup> | *Sec23a*<sup>+/-</sup><br>*Sec23b*<sup>+/b-a</sup> | *Sec23a*<sup>-/-</sup><br>*Sec23b*<sup>+/b-a</sup> | **p-value¥** |
| *Sec23a*<sup>+/-</sup> *Sec23b*<sup>+/b-a</sup> x *Sec23a*<sup>+/-</sup><br>**Expected ratios** | **20%** | **20%** | **20%** | **20%** | **20%** |  |
| Observed at weaning % (n=87) | 16.1% (14) | 19.5% (17) | 28.7% (25) | 35.6% (31) | 0% (0) | <0.0001 |
| Observed at E14.5 % (n=18) | 2 | 2 | 4 | 10 | 0% (0) | 0.011 |
| Observed at E13.5 % (n=41) | % (5) | % (4) | % (22) | % (10) | 0% (0) | <0.0001 |

An intercross between mice heterozygous for the *Sec23b*^*b-a*^ allele generated the expected number of *Sec23b*^*b-a/b-a*^ offspring at weaning (p = 0.68) (Table 1b). In WT mice, SEC23B is the predominantly-expressed SEC23 paralog in the pancreas (Fig 3d). Consistent with substitution of SEC23A coding sequence for SEC23B, no SEC23B immunoreactive material was detected in pancreas tissues of *Sec23b*^*b-a/b-a*^ mice, with high levels of SEC23A protein observed (Fig. 3d), in contrast to WT mice, which exhibit the opposite pattern. SEC23B was also undetectable in *Sec23b*^*b-a/b-a*^ liver and lung tissues (Fig 3d).

*Sec23b*^*b-a/b-a*^ mice exhibited normal growth (Fig 3e, f) and fertility, normal gross appearance, no abnormalities on histopathologic evaluation (Extended Data Fig. 6-7), and normal overall survival (up to 1.5 years of follow-up) (Fig. 3g). In contrast to the pancreatic degeneration and distended ER observed in *Sec23b*^*−/−*^ mice^10,12^, pancreas tissue from *Sec23b*^*b-a/b-a*^ mice exhibited normal weight (Fig. 3h) and histology (Fig. 3i), indistinguishable from pancreas of WT littermate controls. The *Sec23b*^*b-a/b-a*^ pancreatic ER also appeared normal by transmission electron microscopy (Fig. 3j), with intact acinar cell zymogen granules (Fig. 3k), in contrast to the appearance in *Sec23b*^*−/−*^ mice (Fig. 3j, k). Other secretory tissues that were perturbed in SEC23B-deficient mice^10^ also appeared entirely normal histologically in *Sec23b*^*b-a/b-a*^ mice (Extended Data Fig. 6). Taken together these findings demonstrate that the SEC23A protein rescues both the lethality and the massive pancreatic degeneration of SEC23B-deficient mice, when expressed under the endogenous regulatory elements of the *Sec23b* gene. Although mice homozygous for the latter genetically engineered *Sec23b* locus (expressing only SEC23A and no intact SEC23B protein) appear entirely normal and indistinguishable from wild-type controls, a subtle phenotypic difference or key contribution from the residual SEC23B N-terminal tail retained in the SEC23B-A protein (see Fig. 3a) cannot be excluded. Nevertheless, these data demonstrate complete (or near complete) overlap in function between the SEC23A and SEC23B proteins, despite their origin via an ancient gene duplication preceding the origin of vertebrates. As expected, the *Sec23b*^*b-a*^ allele fails to rescue the lethality of mice with germline SEC23A deficiency (Table 1d).

Although the above data suggest functional equivalence for SEC23A and SEC23B in yeast, ZF, and mice, the disparate phenotypes of SEC23B deficiency in humans and mice remain unexplained. To address this question, the relative expression of *SEC23B*:*SEC23A* in WT tissues from both humans and mice was determined by qRT-PCR. This ratio was higher in human BM (9.7) compared to pancreas (1.8), but was higher in mouse pancreas (12.7) compared to BM (2.6), consistent with a greater dependence on SEC23B expression in human hematopoiesis and in the murine pancreas. Taken together, these results suggest that the disparate human/mouse phenotypes are due to an evolutionary switch in the SEC23A/B expression programs between mouse and human tissues.

We previously demonstrated that in contrast to humans, erythroid-specific SEC23B deficiency in mice does not result in anemia or an erythroid phenotype^12^. To test for a role for SEC23A in hematopoiesis in light of the above data, we examined mice with erythroid-specific (*EpoR*-Cre) SEC23A-deficiency, with no erythroid phenotype observed (Extended Data Table 3 and Extended Data Fig. 8 a-g). In contast, mice with combined erythroid SEC23A/B deficiency die at ~E12.5 (Extended Data Table 4) exhibiting reduced size and appeared pale compared to their wild-type litter mate controls (Extended Data Fig 8 h, i), consistent with the requirement for a threshold level of SEC23 expression in the erythroid compartment.

Gene duplications are frequent evolutionary events, with the duplicated copies most commonly accumulating loss of function mutations with subsequent disappearance from the genome^13-15^. Less frequently, paralogous gene copies become fixed either through neofunctionalization (acquisition of a new function by one of the paralogous gene copies), or subfunctionalization (the division of the functions of the ancestral gene between the paralogous copies). The latter process can occur either at the levels of protein function or gene expression. A recent large scale analysis of tissue specific gene expression data suggested that subfunctionalization of expression evolves slowly and is rare among recently duplicated genes pairs, though a more common mechanism among duplicated transcription factors^16^. Consistent with this model, functional equivalence for a number of transcription factor paralogs has been demonstrated in mice by gene replacement^17-20^, although in many cases this overlap is incomplete, resulting in only partial rescue of a null phenotype by the paralogous protein coding sequence^17,20^.

The genes encoding a number of components of the COPII machinery have been duplicated during vertebrate evolution, with evidence for unique protein function for the corresponding paralogous proteins. In the case of SEC23, the unique and highly specific phenotypes of SEC23A and SEC23B deficiency in humans and mice suggested unique functions for each of these paralogous proteins, with evidence for possible paralog specific contributions to collagen^7^ and EGFR^6^ transport, as well as a potential role for SEC23B in oncogenesisis^21^. In contrast, our evidence for functional equivalence between the SEC23A and SEC23B proteins suggests that the evolutionary subfunctionalization of these 2 genes occurred at the level of gene expression program rather than protein function, with select cell types manifesting greater dependence on expression of one or the other paralog. Further shifts in the Sec23a and Sec23b expression programs during mammalian evolution could account for the disparate phenotypes of SEC23A and SEC23B deficiency in mice and humans. Consistent with this hypothesis, human SEC23B is the predominantly expressed paralog in the bone marrow, potentially explaining the bone marrow restricted phenotype of CDAII, while SEC23A is the more strongly expressed paralog in calvarial osteoblasts^22^, consistent with the CLSD phenotype. In contrast, SEC23B is the predominant paralog in the mouse pancreas, consistent with the pancreatic phenotype observed in murine SEC23B deficiency^8-10^.

Similar subfunctionalization at the level of gene expression might also explain highly tissue restricted phenotypes for other genetic diseases due to loss-of-function in paralogous genes encoding other COPII subunits or other core components of the cellular machinery. This paradigm might also explain other examples of highly discordant phenotypes observed between animal models and humans for various other diseases^23^.

Finally, the apparent complete rescue of the SEC23B-deficient phenotype in mice by activating SEC23A expression in SEC23B-dependent tissues suggests that therapeutic strategies to increase the expression level of SEC23A in human hematopoietic precursor cells could ameliorate the CDAII phenotype. Such approaches could include delivery of modified CRISPR/Cas9/transcription factor constructs designed to specifically activate the SEC23A gene^24,25^. If successful, this paradigm could potentially be applicable to treatment for a number of other human disorders due to mutations in functionally-overlapping paralogous genes.

## Materials and Methods

### Co-immunoprecipitation

FLAG-tagged SEC23A and SEC23B, and RFP-tagged SEC24A, SEC24B, SEC24C, and SEC24D were cloned into the PED plasmid. Expression vectors were transfected into COS cells using FUGENE HD (Promega) per the manufacturer’s instructions. FLAG co-immunoprecipitation with anti-FLAG M2 beads (Sigma) was performed as per manufacturer’s instructions.

### Generation of stable cell lines

The coding sequences of human *SEC23A* and *SEC23B* were cloned into the pENTR vector (K2400-20, Invitrogen) per the manufacturer’s instructions and then into the pcDNA5-pDEST-BirA-Flag-C-term vector by Gateway cloning. Bait proteins (SEC23A or SEC23B) were stably expressed in T-REx HEK293 cells as previously described^26^. Stable T-REx HEK293 transformants were selected with 200 µg/ml hygromycin. For “BioID” (Proximity dependent biotinylation) experiments, cells were treated with tetracycline (1 µg/ml) to induce gene expression and with biotin (50 µM) for 24 hours and were subsequently harvested for analysis as previously described^26^.

### Affinity purification and BioID mass spectrometry (MS)

Cell lysates were prepared and biotinylated proteins were captured on streptavidin beads and subjected to trypsin digest as previously described^26^. Tryptic peptides were suspended in 5% formic acid and stored at -80°C until analyzed by MS. Tryptic peptides were analyzed on a TripleTOF 600 instrument as previously described^26^. Two biological replicates were analyzed for each sample and for controls, and data analysis was performed as follows. Raw data was converted to mzXML and searched using the X! Tandem search engine. Xxxx database was appended with equal number of decoy sequences, xxxx precursor tolerance and xxx fragment tolerance. The search results were further processed using PeptideProphet to generate a list of peptides, which were then assembled to protein lists using ProteinProphet. The resulting protein lists were then processed using Prohits (default parameters) to generate a spectral count matrix. The spectral count matrix was formatted according to CRAPome/REPRINT specifications (manually) and uploaded into re-print-apms.org. Protein interactions were scored using two enrichment scoring approaches, empirical fold change score (FC) and SAINT (SP). A list of high confidence interactions was generated by applying the following filters: FC > xxx; SP > xxx). Finally, high scoring interactions were visualized as a network (using the network generation utility of REPRINT). Raw abundances of filtered interactions in each replicate were plotted against other to evaluate correlation of prey abundances….

### Sec23 complementation in yeast

The coding sequences of the mouse and human *SEC23* paralogs, (as well as control yeast *SEC23*) were cloned into the yeast expression vector pRS416, with a 3’ sequence encoding an HA tag. The sequence integrity of all constructs was confirmed by Sanger sequencing. Vectors with the following promoters were used: the *ZEO1* constitutive promoter, the *CUP1* copper-inducible promoter, and the *GAL1* galactose-inducible promoter, resulting in a range of expression of the SEC23 proteins. The latter vectors were transduced into a temperature sensitive *sec23-1* yeast mutant in the BY4741 background^27,28^, which expresses the functional endogenous yeast Sec23 protein at the permissive temperature (24°C) but not at the restrictive temperature (37°C).

### *Sec23b* Morpholino knock-down in Zebrafish (ZF)

One-cell stage ZF embryos were injected with a *Sec23b* splice targeting morpholino (Mo; Extended Data Table 1) diluted in injection buffer (experimental group) or with injection buffer only (control group), as previously described. Blood was collected from 4- to 5-day-old embryos, and cytospins were prepared and examined under light microscopy for percentage of bi-nucleated erythroid cells.

### Generation of SEC23B-deficient ZF using CRISPR/Cas9

The *Sec23b* locus was identified in the ZF genomic sequence assembly^29^. A guide RNA (gRNA) targeting the ZF *Sec23b* exon 5 sequence (ggtggacacatgtctggagg) was cloned into the pDR274 vector as previously described^30^. gRNA was then *in vitro* transcribed using the T7 Quick High Yield RNA Synthesis kit (NEB) and purified using the RNA Clean and Concentrator kit (Zymo Research). gRNA was co-injected with Cas9 RNA (generated as previously described^30^) into single-cell zebrafish embryos. ZF were genotyped for indels with primers ZF *Sec23b* CRISPR F and ZF *Sec23b* CRISPR R (Extended Data Table 1) spanning the expected CRISPR cut site. ZF heterozygous for a 53 base-pair deletion in exon 5 of *Sec23b* (*Sec23b*^*+/−*^) were generated and crossed to wild-type (WT) ZF expressing dsRed in erythrocytes under the control of the *Gata1* promoter^31^. *Sec23b*^*+/−*^ ZF were intercrossed to generate Sec23b-deficient ZF (*Sec23b*^*−/−*^). Blood collected from WT, *Sec23b*^*+/−*^, and *Sec23b*^*−/−*^ ZF was stained with the HEMA 3 kit (Fisher) and evaluated under light microscopy by 2 independent investigators blinded to the ZF genotype.

### Generation of transgenic ZF expressing SEC23A under the control of a ubiquitin promoter

The ZF *Sec23a* cDNA was cloned downstream of the ubiquitin promoter into the JS218 vector^32^ (SEC23A-GFP). The correct *Sec23a* sequence was confirmed by Sanger sequencing. A p2A sequence separates the *Sec23a* cDNA and the GFP coding sequence, resulting in ubiquitous *Sec23a* and *GFP* expression from the same mRNA molecule (Extended Data Fig. 3).

SEC23A-GFP or JS218 (in which GFP alone is expressed ubiquitously) were co-injected with transposase mRNA as previously described^33^ into one-cell stage embryos generated from *Sec23b*^*+/−*^ ZF intercrosses to generate transgenic ZF ubiquitously expressing SEC23A and GFP or GFP alone. The latter ZF were genotyped with primers ZF *Sec23b* CRISPR F and ZF *Sec23b* CRISPR R (listed in Extended Data Table 1).

### Cloning the *Sec23b-a* dRMCE cassette into pUC19

The *Sec23b-a* cassette was generated by assembling the following sequences from the 5’ to 3’ direction: an FRT sequence, the endogenous *Sec23b* intron 4 splice acceptor sequence, *Sec23a* from C^367^ to A^2298^ of the cDNA coding sequence (encoding murine SEC23A starting at Arg123), and the SV40 polyA sequence present in the *Sec23b*^*cgt*^ allele^8,34^. The above *Sec23b-a* cassette was inserted into pUC19 at the *HindIII* and *SalI* restriction sites, producing pUC19-Sec23b-a (1) (Extended Data Fig. 4) and a puromycin resistance cassette flanked by *rox* sites was inserted 3’ of the SV40 polyA sequence using *PmeI* and *AscI* restriction sites added to the vector by PCR, resulting in pUC19-Sec23b-a(2) (Fig. 3a), with sequence integrity confirmed by Sanger sequencing.

### Plasmid purification and microinjections

The pDIRE plasmid driving expression of both FLPo and iCRE was obtained from Addgene (#26745). The pCAGGS-FLPo and pCAGGS-iCre plasmids, which contain the CAG promotor/enhancer driving the expression of FLPo and iCre recombinase respectively, were generated as previously described^35^. Plasmids were purified using the Qiagen EndoFree Plasmid Maxi Kit (Qiagen, #12362) or the Machery-Nagel NucleoBond Xtra Maxi EF kit (Machery-Nagel, #740424.10), per the manufacturers’ instructions.

Zygotes generated from the *in vitro* fertilization of C57BL/6J oocytes with sperm from *Sec23b*^*+/−*^ male mice^12^, were co-injected with pUC19-Sec23b-a(1) and pDIRE plasmids and then implanted in pseudopregnant foster mothers. Pups were genotyped at 2 weeks of age. All microinjections were performed at the University of Michigan Transgenic Animal Model Core.

### Transient electroporation of ES cells

The ES cell clone EPD0237_3_G06 heterozygous for the *Sec23b*^*cgt*^ allele^12^ was obtained from the European Conditional Mouse Mutagenesis Program^34^ and co-electroporated with pUC19-Sec23b-a(2) and either pDIRE or a combination of pCAGGS-FLPo and pCAGGS-iCre, as previously described^35^. One week post-electroporation, individual ES cells were plated in 96-well plates. Cells were expanded and plated in triplicates for DNA analysis, frozen stocks, and puromycin selection. Following puromycin selection, 288 single cell ES clones were evaluated demonstrating correct targeted insertion of the dRMCE replacement vector in 13/288 independent clones.

### Generation of *Sec23b*^*+/b-a*^ mice

Three independent EPD0237_3_G06 ES cell clones correctly targeted with pUC19-Sec23b-a(2) were cultured, expanded, and then microinjected into albino C57BL6 blastocysts to generate *Sec23b*^*+/b-a*^ mice (Fig. 3a). The chimeric progeny were crossed to B6(Cg)-Tyr^c-2J^/J mice (Jackson laboratory stock #000058) and germline transmission was confirmed by genotyping the progeny for the *Sec23b*^*+/b-a*^ allele. The *Sec23b*^*+/b-a*^ allele was maintained on a C57BL/6J background.

### *Sec23a* and combined *Sec23a*/*Sec23b* deletion in the erythroid and pan-hematopoietic compartments

*Sec23a*^*fl*^ mice^8,12^, in which exon 3 is flanked by *loxP* sites, were crossed to mice expressing Cre recombinase under control of the erythropoietin receptor promoter (*EpoR*-Cre mice) (generous gift from Dr. Ursula Klingmüller^36^) to generate mice with erythroid-specific SEC23A deficiency. Similarly, mice were generated with combined erythroid deletion for *Sec23a* and *Sec23b* (Extended Data Table 3).

### Mouse genotyping

Genotyping for the *Sec23b* and *Sec23a* floxed and null alleles as well as for the *EpoR*-Cre allele was performed as previously described^8,12^. Genotyping of the *Sec23b*^*b-a*^ allele was done on genomic DNA extracted from tail biopsies using primers RMCE F2 and RMCE R2, which detect the dRMCE replacement construct. These primers are located in *Sec23a* exons 10 and 14, respectively, and therefore should not yield a detectable product from the wild type *Sec23a* allele. To identify potential random insertions of the dRMCE allele, PCR genotyping was also performed with primers RMCE F1 and RMCE R1 to confirm correct insertion of the Sec23b-a(1) and Sec23b-a(2) alleles at the 5’ end, and with primers RMCE F3 and RMCE R3 to confirm correct placement of the Sec23b-a(2) allele at the 3’ end. All mice with a visible PCR product with primers RMCE F2 and RMCE R2 also had PCR products with primers RMCE F1-RMCE R1 and primers RMCE F3-RMCE R3, effectively excluding random insertion.

Genotyping for the *Sec23b*^*b-a*^ allele was also performed by RT-PCR on RNA isolated from mouse tail biopsies, using a three-primer competitive PCR assay, with a forward primer (RMCE A) located in exon 2 of *Sec23b* and 2 reverse primers, one located in the *Sec23a* cDNA (RMCE B) and one located in exon 5 of *Sec23b* (RMCE C). This competitive PCR assay should generate a 282-bp band from the WT allele and a 311-bp band from the *Sec23b*^*b-a*^ allele (Fig. 3c), which were resolved on a 2% weight/volume agarose gel. The location of the primers is indicated in Fig. 3a and Extended Data Fig. 4 and the primer sequences are listed in Extended Data Table 1.

### RT-PCR

RNA was isolated from tissues using the RNeasy Mini kit (Qiagen), and reverse transcription (RT) was performed using the SuperScript first strand synthesis system for RT-PCR (Invitrogen) with random primers.

### Western blot and antibodies

Total cell lysates were prepared from pancreas tissues and bone marrows, and western blots were performed as previously described^12^. Rabbit anti-SEC23A and anti-SEC23B antibodies were generated as previously described^12^ and mouse anti-GAPDH and anti-RAL-A antibodies were obtained from Millipore and BD Biosciences, respectively. Anti-FLAG, anti-RFP, and anti-HA antibodies were obtained from Cell Signaling Technology, abcam, and Sigma, respectively.

### Complete blood counts

Peripheral blood was collected from the retro-orbital venous sinuses of anesthetized mice as previously described^8^, and complete blood counts were performed using the Advia120 whole blood analyzer (Siemens), as previously described^12^.

### FACS sorting

Mouse bone marrows were harvested, stained for erythroid cells, and sorted as previously described^12^.

### Animal care

Mice were housed at the University of Michigan and all animal care and procedures were in accordance with the Principles of Laboratory and Animal Care, established by the National Society of Medical Research. Mice had free access to water and food and were kept in cages in a 12-hour light/dark cycle. All animal protocols used in this study were approved by the University of Michigan Committee on Use and Care of Animals.

### Histology, cytology, and electron microscopy

For histological analysis, mouse tissues were harvested and immediately fixed in 4% paraformaldehyde or aqueous buffered zinc formalin (Z-fix, Anatech). Tissues were then routinely processed, paraffin-embedded, sectioned, and stained with hematoxylin and eosin (H&E). Bone marrow smears were prepared using a brush technique, air dried and stained with modified Romanowsky staining (Diff-Quik) technique. For electron microscopy, mouse pancreas tissues were prepared as previously described^8,9^ and sections were examined on a Philips CM100 electron microscope. Images were recorded as previously described^8,9^.

### Statistical analysis

The statistical significance of deviation from the expected Mendelian ratio of mouse crosses was assessed using the Chi-square test. The statistical significance of the differences between parameters among different genotype groups was assessed using the student’s t-test. The difference in overall survival between mice of distinct genotypes was assessed using the Kaplan Meier method.

## Acknowledgments

This work was supported by National Institute of Health Grants R01 HL039693 and P01-HL057346 (DG), K08 HL128794 (RK), R01 GM053396 (DJK), R01 HL 124232 (JAS), T32 HL007622 (ACW). Rami Khoriaty and Jordan Shavit are recipients of the American Society of Hematology Scholar Award. David Ginsburg is a Howard Hughes Medical Institute investigator. Angela Weyand was supported by a National Hemophilia Foundation-Shire Clinical Fellowship Award. The authors would like to acknowledge Elizabeth Hughes, Corey Ziebell, Galina Gavrilina, Wanda Filipiak, Keith Childs, and Debora VanHeyningen for microinjections and preparation of ES cell-mouse chimeras. Support for the Transgenic Animal Model Core of the University of Michigan’s Biomedical Research Core Facilities was provided by the University of Michigan Multipurpose Arthritis Center (NIH grant AR20557) and the University of Michigan Cancer Center (NIH grant P30 CA046592). The authors would like to acknowledge Wendy Rosebury-Smith and Kathy Toy of the In Vivo Animal Core for their necropsy and histology expertise.

